# Java sparrow song conforms to Menzerath’s Law but not Zipf’s Law of Abbreviation

**DOI:** 10.1101/2023.12.13.571437

**Authors:** Rebecca N Lewis, Anthony Kwong, Masayo Soma, Selvino R de Kort, R Tucker Gilman

## Abstract

Linguistic laws, which describe common statistical patterns in human language, are often studied in animal communication systems. Despite good evidence for adherence to these laws in non-human primate communication, there is still a paucity of data on other taxa. Birdsong is often used as a model for human language development due to similarities in vocal learning processes. Investigating whether birdsong also adheres to linguistic laws may provide further value or help determine the limitations of such comparisons. Here, we looked for patterns consistent with Menzerath’s law, the tendency for longer sequences to be made up of smaller constituent parts, and Zipf’s Law of Abbreviation, the tendency for more frequently used elements to be shorter in length, in the songs of Java sparrows (*Padda oryzivora*). We found that songs conformed to Menzerath’s law at both the population and individual level; longer sequences were made up of shorter notes. This pattern was achieved through note type selection rather than the shortening of existing note types. Adherence to Menzerath’s law may reflect species-specific constraints on song durations. We found no evidence to support adherence to Zipf’s Law of Abbreviation, either at population or individual level; more frequently used notes were not shorter in length. However, as repertoire size in Java sparrows is small, we expect our power to detect Zipf’s Law of Abbreviation to be weak. Whilst there may be some limitations in applying linguistic laws derived from human language to birdsong, especially Zipf’s Law of Abbreviation, these laws may still provide insights into animal and human communication systems.

## Introduction

Birdsong and spoken language share many similarities (Berwick et al., 2011; Bolhuis et al., 2010; Hyland Bruno et al., 2021). Both are made up of acoustic sequences, and there is overlap in the behavioural, neural, and genetic bases of vocal learning and production in songbirds and humans (Berwick et al., 2011, 2012; Bolhuis et al., 2010; Hyland Bruno et al., 2021). Due to these similarities, birdsong is often used in comparative studies examining the evolution and development of human language (e.g., Berwick et al., 2011). However, there are also differences between birdsong and human language. In particular, the close relationship between structure and meaning in human language is thought not to exist in other animals (Bolhuis et al., 2010). Testing the limitations of the parallels between human and animal communication may provide insight in the universal laws affecting acoustic communication.

Linguistic laws describe common statistical patterns in human language, where they have been studied extensively (Semple et al., 2022). More recently, patterns consistent with linguistic laws have also been explored in animal communication systems (Semple et al., 2022). Two laws that often receive attention are Menzerath’s law, the tendency for longer sequences to be made up of smaller constituent parts, and Zipf’s Law of Abbreviation (ZLA), the tendency for more frequently used elements to be shorter in length. In human languages, length is often equated with the number of letters in a word, whereas for animal communication the duration (e.g., in seconds) of the element is usually measured (REFS?). These laws are especially well studied in non-human primates (Bezerra et al., 2011; Clink et al., 2020; Clink and Lau, 2020; Gustison et al., 2016; Huang et al., 2020; Semple et al., 2010; Valente et al., 2021; Watson et al., 2020) and some other mammal taxa (Demartsev et al., 2019; Ferrer-i-Cancho et al., 2013; Youngblood, 2025). Studies examining the presence of linguistic laws in bird communication are still relatively limited (Favaro et al., 2020a; Ferrer-i-Cancho et al., 2013; Hailman et al., 1985; James et al., 2021a; Wascher & Youngblood, 2025; Youngblood, 2024).

Adherence to Menzerath’s law has been widely reported in non-human primates (Clink et al., 2020a; Gustison et al., 2016; Heesen et al., 2019; Huang et al., 2020a; Valente et al., 2021). Despite this, adherence to the law is not universal, and may depend on the subset of vocalizations examined; in mountain gorillas, Menzerath’s law was apparent when single-unit sequences were included in analyses, but not when they were excluded (Watson et al., 2020). Patterns consistent with Menzerath’s law have also been found in a range of songbird species, both at the repeat phrase and whole song level (James et al., 2021b; Youngblood, 2024). However, the mechanisms by which these patterns arise have not been explored in detail in songbird species with complex vocalizations. In the African penguin (*Spheniscuc demersus*), patterns consistent with Menzerath’s law are achieved through a combination of note choice and reducing the length of specific note types (Favaro et al., 2020b). However, the vocal sequences of penguins are relatively simple compared to birdsong, so determining if the same mechanisms operate in more complex avian vocal sequences could be beneficial for comparative studies examining the evolution of language.

ZLA has been reported in a range of mammal taxa (Demartsev et al., 2019; Ferrer-i-Cancho & Hernández-Fernández, 2013; Semple et al., 2010b; Valente et al., 2021; Youngblood, 2025), but is not universally found across studies (Bezerra et al., 2011; Clink et al., 2020b; Youngblood, 2025). As with Menzerath’s law, adherence to ZLA may depend on the subset of the repertoire studied; in common marmosets (*Callithrix jacchus*), long-distance calls did not adhere to ZLA, whereas short-range signals were consistent with the expected pattern (Ferrer-i-Cancho & Hernández-Fernández, 2013). Adherence to ZLA may also depend on the unit of analysis. Across gibbon species, ZLA has been reported at the note level, with shorter duration notes being used more frequently in loud morning calls for the cao vit (*Nomascus nasutus*) and western black-crested (*Nomascus concolor*) gibbons (Huang et al., 2020b), but not at the phrase level in male solos of the Mueller’s gibbon (*Hylobates muelleri*) (Clink et al., 2020b). ZLA is not commonly studied in birds, although a small number of species have been examined to date (Favaro et al., 2020b; Ferrer-i-Cancho et al., 2013; Gilman et al., 2025; Hailman et al., 1985; Youngblood, 2024). In African penguin calls, shorter duration note types were used more frequently than longer ones, a pattern consistent with ZLA (Favaro et al., 2020b). However, these calls are relatively simple and contain only a few note types. In the common raven (*Corvus corax*), researchers found no significant evidence for ZLA (Ferrer-i-Cancho et al., 2013). More recently, patterns consistent with ZLA have been reported in house finch (*Haemorhous mexicanus*) songs (Youngblood, 2024). Gilman et al. (2025) found only weak evidence for ZLA in individual populations of seven bird species, but a synthetic analysis across species supported the presence of ZLA in birdsong generally. There is, therefore, a need to study more populations of birds with complex note repertoires to fully understand ZLA in birdsong.

Here, we looked for patterns consistent with Menzerath’s law and ZLA in Java sparrow (*Padda oryzivora*) song (Lewis et al 2023). Each individual male Java sparrow produces a single song type that is repeatable but varies across renditions (Ota & Soma, 2014). This variability in note usage both among and within individuals makes the Java sparrow a good candidate for examining linguistic laws in birdsong. Our goals were i) to assess whether Java sparrow song adheres to Menzerath’s law, and, if so, to determine whether birds achieve this by choosing shorter notes or by producing shorter versions of the same notes in longer songs; and ii) to determine whether Java sparrow song adheres to ZLA at the individual and population levels.

## Methods

### Study species

The Java sparrow (*Padda oryzivora*) is an oscine passerine and member of the family Estrildidae (Restall, 1996). Songs in this species are used in courtship but not for territory defence. Each male learns to produce a single song with 2-8 note types from his social father (Lewis et al., 2021, 2023; Soma, 2011) and produces his song with some variation among renditions (Ota & Soma, 2014). Learning fidelity is high, with sons’ songs resembling their fathers’ songs in a range of spectral and temporal parameters (Lewis et al., 2021, 2023).

### Dataset

To investigate the presence of linguistic laws in Java sparrow song, we used the dataset from Lewis et al. (2023), which contains 22,966 notes across 676 undirected songs from 73 individuals (see Figure 1 for examples). Birds in this dataset belonged to 10 social lineages. Each social lineage included a single founder, all the males that learned their songs from that founder (i.e., his social sons), the social sons of those birds, and so on for up to five generations. Thus, songs within social lineages were not independent. Some birds (n=6) were reared in the presence of subtutors (an additional adult male that was not their social or genetic father). Spectrogram inspection suggests that subtutored birds generally produced songs that resembled their social fathers’ song. Therefore, subtutored individuals were classed as belonging to their social fathers’ lineage for this study. Songs were defined as series of notes with inter-note intervals <1 s (Kagawa & Soma, 2013; Lewis et al., 2021). Songs were manually segmented into individual notes using the software Koe (Fukuzawa et al., 2020) and classified into 16 note types, with high inter-observer repeatability (Lewis et al., 2021).

**Figure 1:**
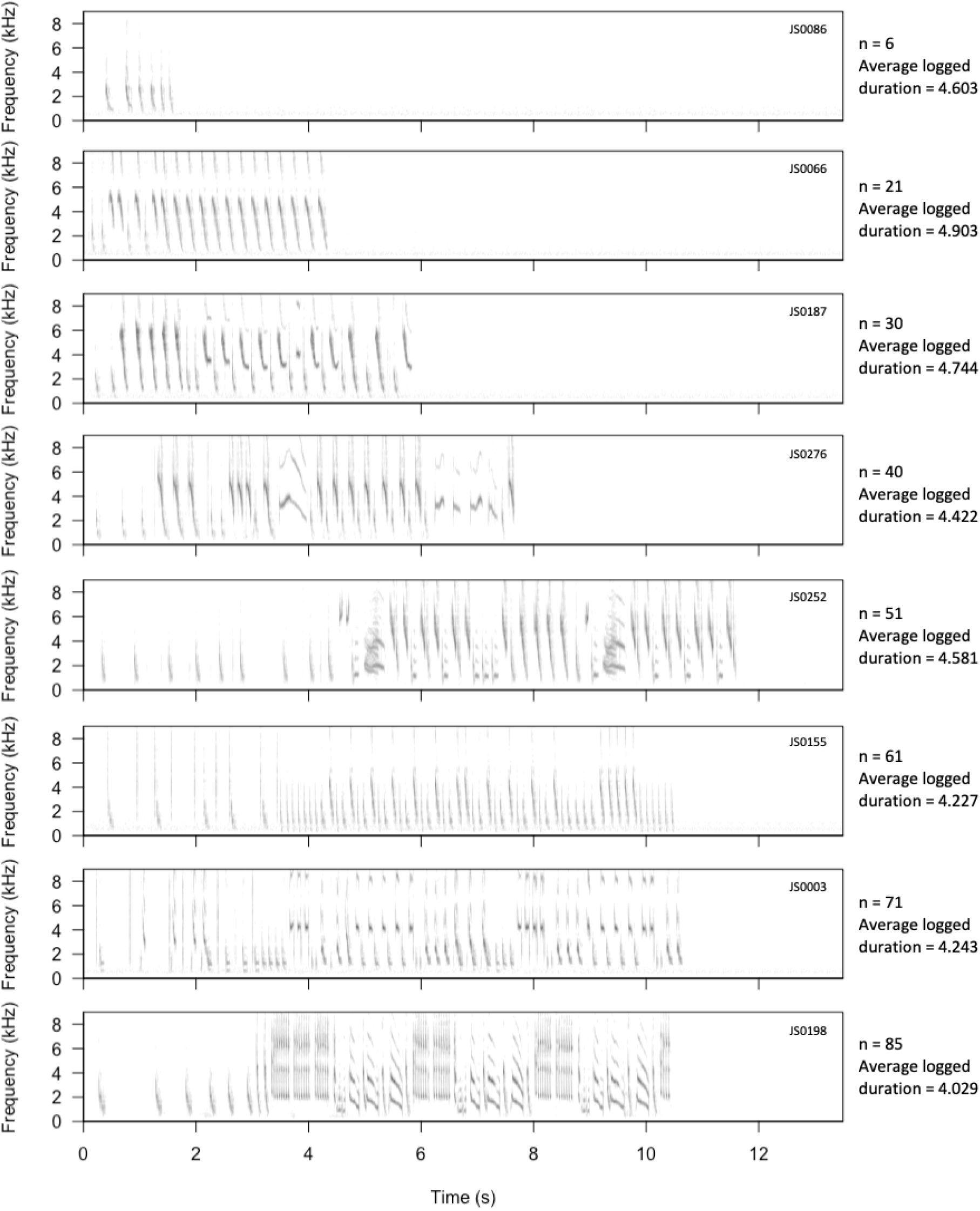
Example spectrograms of songs from 8 individual Java sparrows from different social lineages. Number of notes and the average log-transformed durations of notes for each song are indicated next to spectrograms. Spectrograms were created using Seewave (Sueur et al., 2008) (sample rate = 44.1kHz, window length = 512, overlap = 90%).

Java sparrow songs frequently begin with a single note or small sequence of notes repeated several times with uncharacteristically long gaps (>0.25 s but <1 s) between repetitions (Lewis et al. 2023). These introductory sequences may represent periods of preparation prior to main songs (as in zebra finches e.g., (Kalra et al., 2021; Rajan & Doupe, 2013; Rao et al., 2019)), but may also be important in song learning and development (Kalra et al., 2021). Therefore, it is difficult to determine whether these notes should be counted as part of the main song sequence. As introductory notes contribute to sequence length (i.e., the number of notes in a song), the presence of introductory notes may affect our analysis of linguistic laws, and in particular, Menzerath’s law. To account for the presence of introductory notes, all analyses concerning Menzerath’s law were conducted twice, once with and once without introductory notes included, to determine if patterns were similar in both cases (Appendix 1). As in Lewis et al. 2023, we considered the bird to have begun its main song as soon as it began a sequence of 5 or more notes with inter-note intervals of <0.25 s, and we considered any notes before the beginning of the main song to be introductions. The full dataset included 20911 notes when song introductions were excluded. Results were qualitatively similar for analyses with and without introductory notes, with minor differences indicated in the ‘Results’ section.

### Data Analysis

#### Menzerath’s Law

We examined adherence to Menzerath’s law by asking i) whether longer songs (i.e., those with more notes) are made up of shorter duration notes in the population as a whole. This is how Menzerath’s law has been studied in other animals (Gustison et al., 2016; Valente et al., 2021). Java sparrow songs are composed of different note types, and note duration depends on both the note type and the singer (Lewis et al., 2021). To examine how Menzerath’s law might arise in Java sparrow song, we asked ii) whether notes of the same type have shorter durations when they appear in longer songs, and iii) whether longer songs are composed of shorter note types. Menzerath’s law could arise at the population level due to differences in song production among birds even if the same law does not hold within individual birds. For example, we would see this if some birds produce only long songs composed of short duration notes and other birds produce only short songs composed of long duration notes. Therefore, we conducted a second set of analyses to ask whether Menzerath’s law holds within individuals. In particular, we asked, iv) whether individual Java sparrows sing shorter duration notes when they produce longer songs, v) whether individuals use shorter duration versions of the same note types in longer songs, and vi) whether individuals use shorter duration note types in longer songs. To our knowledge, Menzerath’s Law has not previously been studied at the level of individuals in birdsong, although the importance of studying Menzerath’s law at the individual level has been advocated for human language (Grzybek & Stadlober, 2007).

To answer the above questions, we fit a series of linear mixed effect models using the lme4 (Bates et al., 2015) and lmerTest packages (Kuznetsovs et al., 2017) in R (ver 4.1.3) (R Core Team, 2022). To address question i) we regressed the log-transformed duration of individual notes on the length of the song (defined as the number of notes in the song) that contained them, and on their log-transformed position in the song. A significant negative relationship between the note duration and song length would be evidence for Menzerath’s law, and a significant relationship between note duration and position in the song would indicate that note duration changed systematically over the course of each song (i.e., notes later in the song are shorter or longer in duration than those at the beginning). Note duration was log-transformed to stabilize variance, as longer notes are inherently more variable than shorter notes. Position in the song was log-transformed, as we expected the effect of position in the song to increase less than linearly with the position, and models with log-transformed position provided lower AICc values. AICc values were not consistently lower for models with log-transformed song length. Therefore, we fit models to untransformed song length fot consistency with other recent work (Youngblood, 2024). To account for the non-independence of note durations within songs, individuals and social lineages, we included random effects of song, individual bird, and social lineage in the model. The random effects of song included a random intercept, to account for the possibility that some songs might have longer notes than others, and a random slope of position in song, to account for the possibility that note length might change differently over the course of different songs. The random effects of bird and social lineage included intercepts, slopes of log-transformed position in the song, and slopes of song length (number of notes in the song). The random intercept accounts for the possibility that different birds or social lineages might sing notes of different durations, the random slope of log-transformed position accounts for the possibility that note duration may change differently over the course of songs in different individuals or lineages, and the random effect of song length accounts for the possibility that song length might affect note length differently in different individuals or lineages. There was no random slope of song length within songs because each song has only one length. The random slopes of song length within individuals and lineages allows that the strength of Menzerath’s law may differ among individuals and lineages.

Adherence to Menzerath’s law at the population level could be achieved in two ways; birds might sing shorter duration versions of the same note types in longer songs (question ii), or birds might use shorter duration note types in longer songs (question iii). To disentangle these two possibilities, we conducted two regressions identical to the one just described, but with different response variables. To address question ii) the response variable was the log-transformed duration for each note in the dataset, mean centred within its note type. Thus, differences in note duration due to the note types are removed, and any patterns that remain must be due to differences in how notes of each type are produced. To address question iii) we replaced each log-transformed note duration in the dataset with the mean log-transformed duration for its note type in the whole dataset. Thus, variability within note types is removed, and any pattern that remains must be due to the note types that birds use in their songs.

Java sparrow songs might adhere to Menzerath’s law across the whole population even if individual birds do not use shorter duration notes in longer songs. To investigate patterns associated with Menzerath’s law within individual birds, we conducted a second set of regressions. In these analyses, we mean-centred the song length within each bird and used the individual mean-centred song length, rather than raw song length as a predictor. Any relationship between individual mean-centred song length and note duration in these analyses cannot be due to differences in song length among individuals, because those have been removed from the predictor, and therefore must be because individual birds use shorter notes in longer songs. Analyses iv to vi are analogous to analyses i to iii, respectively.

#### Zipf’s Law of Abbreviation (ZLA)

We used Kendall’s rank correlation (τ) test to assess the significance of the relationship between the mean log-transformed duration of note types and their frequency of use in our study population. Kendall’s τ has been used to study ZLA in human languages (Bentz & Ferrer-i-Cancho, 2016) and other animal communication systems (Ferrer-i-Cancho et al., 2013; Semple et al., 2010a). Typically, a representative sample of the language (for humans) or vocalizations (for animals) is examined to determine whether the length of individual units is negatively correlated with their frequency of use. However, this simple approach may not transfer directly to birdsong. Individual birds may use highly divergent or even non-overlapping sets of note types, making it difficult to create truly representative samples of note type use in the population. Even if representative samples can be obtained, differences in the use of note types among individuals could create patterns that look like ZLA at the population level but do not arise by the Zipfian mechanism. Zipf (1935) hypothesised that individuals shorten vocal units that they use more frequently to make communication more efficient. If this is true, we should see a negative relationship between the duration and frequency of use of vocal units within as well as among birds. However, if many birds sing only short note types and a few sing only long note types, then the resulting population-level pattern may be consistent with ZLA even if no individual bird uses short note types more frequently than long ones. Another challenge to studying ZLA in birdsong is that birdsong can be highly stereotyped, and is often learned from other members of the population. This means that observations of note type use within and among individuals are not independent, and treating them as independent is likely to result in false positive inferences about adherence to ZLA in populations.

To determine whether our population of Java sparrows adhered to ZLA, we developed a novel test appropriate for bird populations. Briefly, we computed Kendall’s *τ* for note duration and frequency of use within each individual bird, and then computed the mean *τ* over all birds in the population, with each bird weighted for the number of note types in its repertoire. This population-mean *τ* is a measure of the strength of ZLA in the population and serves as a test statistic. To determine whether the population-mean *τ* was larger than we would expect in a random sample from a population without ZLA, we compared it to a null distribution that we computed while maintaining the observed song structure and note similarities among birds. We implemented this method using the R package ZLAvian, which we present in full elsewhere (Gilman et al., 2025).

### Ethical Note

The dataset used in this study was initially collated for use in Lewis et al. (2021) and Lewis et al. (2023). The dataset contains information extracted from archival recordings from a laboratory population of Java sparrows (Padda oryzivora) housed at Hokkaido University. Archive data was collected between 2011 and 2019 under the following approvals: Hokkaido University (Permit numbers: 11-0028, 16-0020) and the University of Manchester AWERB (Permit number: D042).

### Data Availability Statement

Data and code associated with this paper can be found at FigShare https://doi.org/10.48420/24763299. Data and code from Lewis et al. (2023) can be found at Figshare: http://doi.org/10.48420/22550149.

## Results

### Menzerath’s law

We found evidence to support the presence of Menzerath’s law in Java sparrow songs when considering all songs in the dataset (i.e., the population level models i - iii); across birds, as song length (number of notes) increased, notes within the song were shorter in duration (β = -0.004, p=0.025; Table 1; Figure 2). Results were qualitatively similar when considering songs with introductory notes removed, with a trend towards shorter duration notes in long songs (i.e., songs with more notes) (β = -0.003, p=0.084; Appendix 1). At the population level, we did not find that notes significantly changed in duration across the course of the song, although the direction of effect was towards notes being longer when they appeared later in songs (with introductory notes β = 0.058, p=0.121 (Table 1); without introductory notes β = 0.042, p=0.263 (Appendix 1)). When examining the potential drivers of Menzerath’s law in Java sparrow songs, we found that birds did not reduce the durations of note types in long songs (with introductory notes p=0.988 (Table 1); without introductory notes p=0.850 (Appendix 1)). For example, note type ‘A’ in a longer songs was not shorter in duration than note type ‘A’ in a shorter songs. There was also no change over the course of the song (with introductory notes p=0.973 (Table 1); without introductory notes p=0.949 (Appendix 1)); note type ‘A’ later in a song was not shorter in duration than note type ‘A’ earlier in the song. However, we did find evidence that birds used note types with shorter average durations more frequently in longer songs (with introductory notes β = -0.004, p=0.002 (Table 1); without introductory notes β = -0.003, p=0.003 (Appendix 1)). In addition, birds used note types with longer average durations later in songs (β = 0.056, p=0.015; Table 1). Results were qualitatively similar when considering songs with introductory notes removed (β = 0.038, p=0.051; Appendix 1). Overall, these results suggest that adherence to Menzerath’s law is driven by preferential use of short duration note types, rather than by shortening the durations of note types in long songs.

**Table 1:** Results of GLMMs examining the effect of song length (number of notes) and note position in songs on note duration. All durations were log-transformed.

| **Population level** |  |  |  |
| --- | --- | --- | --- |
| *Question* | *Measurement* | *Song length* | *Position in song (logged)* |
| Are longer songs made up of shorter notes? | Log note duration (log(s)) | -0.004<br>$p=0.025$ | 0.058<br>$p=0.121$ |
| Do longer songs contain shorter versions of the same note types? | Deviation from note type mean (log(s)) | $-6.24 \times 10^{-6}$<br>$p=0.988$ | -0.0007<br>$p=0.973$ |
| Are longer songs made up of shorter note types? | Average logged duration of note type (log(s)) | -0.004<br>$p=0.002$ | 0.056<br>$p=0.015$ |
| **Individual level** |  |  |  |
| *Question* | *Measurement* | *Song length* | *Position in song (logged)* |
| Are longer songs made up of shorter notes? | Log note duration (log(s)) | -0.003<br>$p=0.059$ | 0.055<br>$p=0.144$ |
| Do longer songs contain shorter versions of the same note types? | Deviation from note type mean (log(s)) | $-8.41 \times 10^{-6}$<br>$p=0.985$ | -0.0003<br>$p=0.989$ |
| Are longer songs made up of shorter note types? | Average logged duration of note type (log(s)) | -0.003<br>$p=0.009$ | 0.052<br>$p=0.026$ |

**Figure 2:**
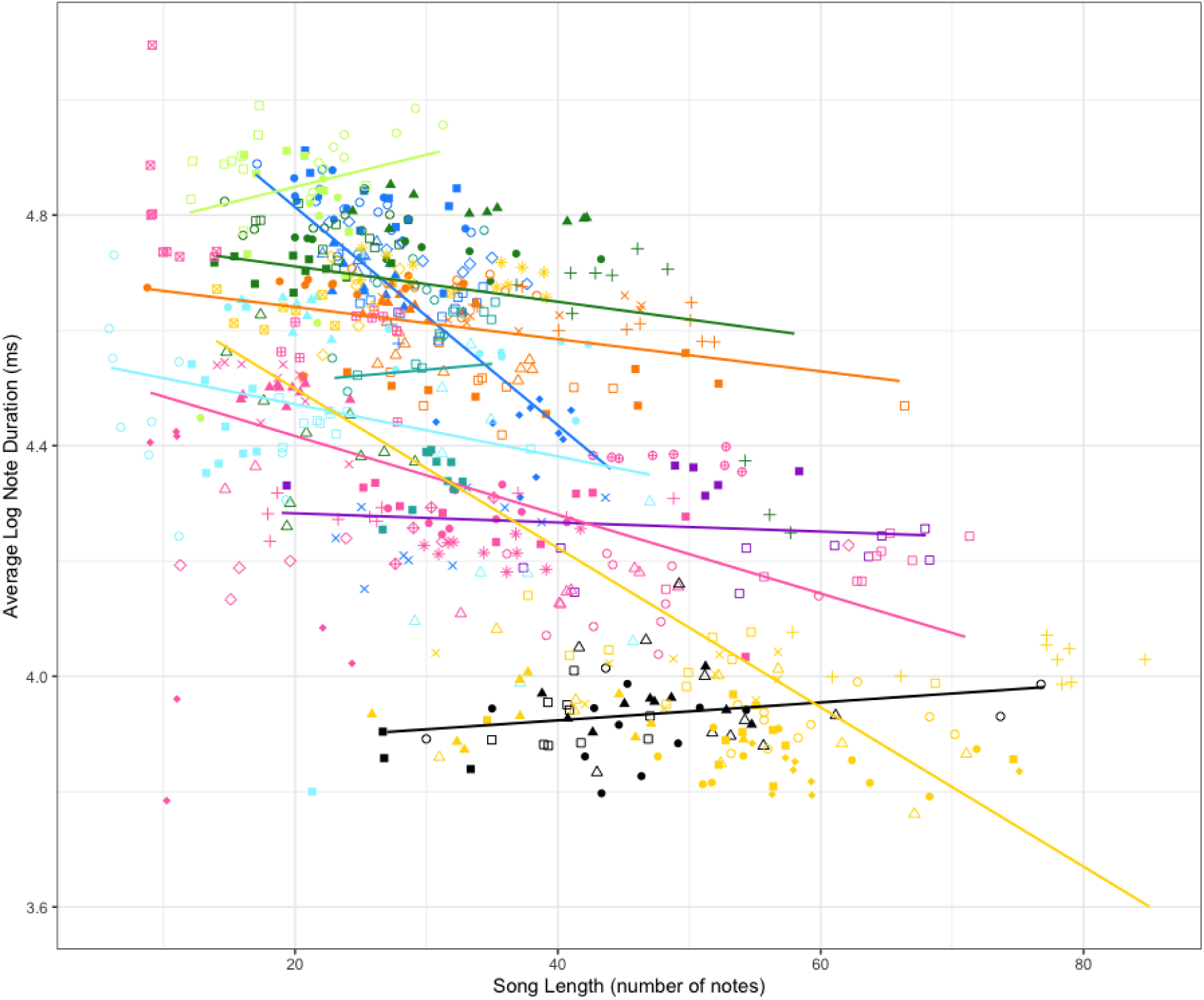
Average log-transformed note duration within a song (ms) plotted against song length (number of notes). Each shape and colour combination represents a single individual. Individuals are coloured based on their social lineage, which relates to song phenotype. Regression lines are calculated for each social lineage and coloured accordingly.

We found weak evidence for Menzerath’s law at the individual level (i.e., models iv-vi); individuals used shorter duration notes when they sang songs with more notes (with introductory notes β = - 0.003, p=0.059 (Table 1); without introductory notes β = -0.002, p=0.084 (Appendix 1)). Again, there was no evidence that note duration became shorter towards the end of the song, and our results suggest a trend in the opposite direction (with introductory notes β = 0.055, p=0.144 (Table 1); without introductory notes β = 0.037, p=0.326 (Appendix 1)). The pattern in individuals was driven by the same factors as the population level findings; birds used shorter duration note types in longer songs (β = -0.003, p=0.009 (Table 1); without introductory notes β = -0.002, p=0.013 (Appendix 1)) and note types with longer average duration later in songs (with introductory notes β = 0.052, p=0.026 (Table 1); without introductory notes β = 0.033, p=0.084 (Appendix 1)). There was no evidence that birds changed the duration of individual note types based on song length (with introductory notes p=0.984 (Table 1); without introductory notes p=0.899 (Appendix 1)) or that notes of a given type were shorter in duration later in songs (with introductory notes p=0.986 (Table 1); without introductory notes p=0.943 (Appendix 1)).

### Zipf’s law of abbreviation (ZLA)

We did not find evidence for ZLA at either the population or individual levels. There was no evidence that shorter duration note types were used more frequently than longer duration note types across all songs in the population (Kendall’s tau; τ=-0.150, p=0.209) or by individual birds (τ=-0.037, p=0.431). The results are qualitatively similar if we exclude note types used by only one bird (population level: τ=-0.282, p=0.090; individual level: τ=-0.030, p=0.441) or if we exclude note types used fewer than 5 times in total (population level: τ=-0.181, p=0.174; individual level: τ=-0.044, p=0.419).

## Discussion

We found evidence that Java sparrow songs adhered to one of the two linguistic laws we studied. Longer songs (i.e., those with more notes) were made up of shorter duration notes across all songs in the population and within the songs of individual birds (Menzerath’s law). This was due to birds using shorter duration note types more frequently in longer songs, rather than shortening the duration individual note types in longer songs. We did not find strong evidence to suggest that note duration changed as the song progressed, although a trend towards longer duration notes later in the song was apparent. However, we found that birds included longer duration note types towards the end of their songs both across the population and within individuals.

Adherence to Menzerath’s law at the level of repeat phrases has been reported previously in Java sparrows (James et al., 2021b), and the same pattern is apparent in this study across full songs. In our study population, songs often contained notes that were not part of repeat phrases, meaning that whole song analysis was more appropriate. Adherence to Menzerath’s law may be due in part to constraints on overall song duration, which may relate to energetic costs, female preferences, or species recognition. In Java sparrows, song length and song tempo are positively correlated; birds that produce songs with more notes sing them at faster rates (Lewis et al, 2023). The current study advances this result by showing that this pattern holds not just for song tempo, but for also for the duration of the notes themselves. At the population level, the social learning of birdsong can create patterns consistent with Menzerath’s law. Each male Java sparrow sings a single song type (Kagawa & Soma, 2013; Lewis et al., 2021; Soma, 2011), and the note types used and the order of note types in each male’s song are learned from his social father (Lewis et al., 2021; Soma, 2011). Therefore, fathers and sons sing similar songs (Lewis et al., 2021; Soma, 2011). This means that correlations between song length and note duration may persist within lineages, with some singing fewer, longer notes and others singing more, faster notes. Our analyses controlled for non-independence of song characteristics within birds and social lineages, so learning alone cannot account for the patterns observed. This is further supported by the presence of Menzerath’s law at the individual level; within an individual, longer songs were also made up of shorter duration notes.

Conformation to Menzerath’s law has been reported in a range of other passerine species (James et al., 2021b), indicating that this pattern may be widespread across the order. However, the methods by which Menzerath’s law arises in birdsong have not been examined in detail. For this population, notes within songs were classified into discrete categories, meaning that we could investigate how patterns consistent with Menzerath’s law were achieved in more detail. Longer songs consisted of note types with shorter average durations, and there was no evidence that the duration of individual note types was compressed in longer songs. Our results indicate that shortening of sequence components in Java sparrows is achieved through note selection alone, rather than through changes to note parameters. Note selection also contributes to adherence to Menzerath’s law in the African penguin (Favaro et al., 2020b), although changes to note parameters also play a role. Mediation of Menzerath’s law through note selection was apparent at both the population and individual level in Java sparrows. As with overall adherence to Menzerath’s law, this suggests that the pattern is not driven solely by a few social lineages with long songs made up of short duration notes, but that individuals also use this mechanism on a by-song basis.

It has been suggested that breathing and performance constraints may result in a negative relationship between sequence position and element duration, i.e., elements towards the end of the sequence are shorter (Gustison et al., 2016). However, we did not find this pattern in Java sparrows. At the population and individual level, we did not find a significant correlation between note position and note duration, although in both cases the direction of effect was towards longer duration notes later in songs. However, at both the population and individual level birds were significantly more likely to use longer duration note types later in their songs. A similar positive correlation between note position and duration has been found in a number of non-human primates, with note duration positively associated with position in the vocal sequence (Clink & Lau, 2020; Huang et al., 2020b; Valente et al., 2021). The presence of longer duration notes later in the song, where a decline in performance may be expected due to energetic constraints, could be an indicator of male quality. In Java sparrows, songs are used during courtship and mate selection. Therefore, indicators of quality, such as the positioning of long duration note types near the ends of songs may be preferred by females. However, it may be that short, fast sections, especially those close to theoretical performance limits (Kagawa & Soma, 2013; Podos, 1997), are more energetically costly, and the use of longer duration notes later in the sequence is indicative of performance constraints in this species. Across a range of species, singing performance declined after sustained singing (Sierro et al., 2023), suggesting that performance limits could play a role in the expression of linguistic laws in this and other bird species. Faithful song learning in the Java sparrow, where birds closely imitate their tutors’ song (Lewis et al., 2021; Soma, 2011), could contribute to the maintenance of songs with longer duration notes at the end of the sequence across generations regardless of performance constraints, especially in songs that are not near to performance limits.

We found no evidence that Java sparrow songs adhered to ZLA, either at the population or individual level. Shorter duration note types were not used significantly more frequently than longer duration note types across the population as a whole or within individual birds’ repertoires. The presence of ZLA has not been well studied in birds. Early studies in the black-capped chickadee found that call bouts with smaller numbers of calls were more frequent than those with more calls (Hailman et al., 1985), but not that calls with fewer notes were more common than those with more notes (Hailman et al., 1985). The frequency of specific note types was not examined. This raises an interesting question regarding the appropriate level of analysis for comparing the acoustic sequences of animals with human language, as it is difficult to equate the various levels of organization (Semple et al., 2022). In Java sparrows, note types are used in different combinations by different individuals, indicating that notes may be the recombinant unit in songs and are an appropriate unit for analysis. A similar approach was used to examine ZLA in the calls of African penguins, with the proportions of individual notes being examined (Favaro et al., 2020b). Penguins’ ecstatic display calls did adhere to ZLA, with shorter syllables making up greater proportions of the calls. However, the calls contained only three syllable types, one of which was inspiratory, making comparisons between these calls and Java sparrow songs difficult. More recently, studies have explored adherence to ZLA in complex bird vocalizations i.e., songs. Whilst some studies report adherence to ZLA (youngblood), a study of ZLA across 11 bird populations found only weak concordance with ZLA for any individual population (Gilman et al., 2025). Although a synthetic analysis found evidence for ZLA more broadly, the negative concordance between duration and frequency of use was several times weaker than that reported between word length and frequency of use in written human languages (Gilman et al., 2025). If patterns associated with ZLA are weak in birdsong, then species, including the Java sparrow, may not use enough individual note types to infer statistical significance. Indeed, while evidence for ZLA in Java sparrows was not significant, the estimated concordance between note duration and frequency of use was -0.037, which is within the range of values reported by Gilman et al. (2025) for the populations they studied. It may be necessary to study songs of many different populations and species to build a more general understanding of ZLA across birds, and this study contributes to that effort.

If ZLA is weaker in birdsong than in human languages, as our results suggest, it may be due to a range of phenomena that select against compression of vocal signals in birdsong (Ferrer-i-Cancho et al., 2013). Java sparrow songs are used in courtship and mate choice, and so may be under different selective pressures compared to other parts of the vocal repertoire. In this case, compression and signal efficiency might not be an advantage, as songs can be honest indicators used to advertise male quality (Gil & Gahr, 2002). This may act against compression in several ways. If longer duration notes (or the use of multiple long notes) are signals of quality, longer duration notes may be overrepresented in songs due to their effect on female preference. Similarly, certain note types with particularly attractive qualities, such as ‘sexy syllables’ in canaries (Leitner et al., 2001; Vallet & Kreutzer, 1995), may be over-represented. Faithful social learning of song features, as is apparent in Java sparrows (Lewis et al., 2021, 2023), may also act against compression of vocal signals by preventing compression of note types and changes in the proportion of different note types within songs. Unlike human language, the individual ‘meaning’ of notes and songs, or how these are perceived by the receiver, is not known. It is possible that signal compression affects perception by the receiver. In the closely related Bengalese finch, female-directed songs are more consistent in terms of acoustic structure of individual notes and sequence structure (Sakata et al., 2008). This suggests that even small changes to notes within songs may be salient to females. Java sparrows produce a range of call types in addition to songs (Restall, 1996). Calls often have specific meaning (Marler, 2004), and so this subset of the repertoire may exhibit different patterns relating to compression. In the common marmoset, long distance calls, which may be more affected by attenuation and signal degradation, did not conform to ZLA, whereas short range calls were consistent with the expected pattern (Ferrer-i-Cancho & Hernández-Fernández, 2013).

Overall, we find that Java sparrow songs adhere to Menzerath’s law - longer sequences are made up of shorter constituent parts – but we found no strong evidence for ZLA – more frequently used note types do not have significantly shorter durations. It is likely that these laws reflect different components and constraints on birdsong. Adherence to Menzerath’s law may relate to a constraint on song duration, where songs with more notes necessarily have a faster tempo. However, ZLA is more closely related to meaning and information transfer, which is difficult to attribute to birdsong. It has been suggested that some aspects of birdsong may be more suited to comparisons with human music, rather than language, due to their repetitive nature, aspects of performance, and lack of semantic meaning (Rothenberg et al., 2014). The case of meaning and information transfer may be a case where comparisons to music are more appropriate than comparisons to language. Our findings suggest that although there may be some limitations in applying linguistic laws designed for human language to birdsong, they may nonetheless provide insight into communication universals and warrant further investigation.

## Author Contributions

Conceptualization: RL, AK, TG; Data curation: RL, MS; Formal analysis: RL, AK, TG; Funding acquisition; RL, MS, TG; Investigation – RL, MS, SdK, TG; Methodology: RL, AK, TG; Resources: MS; Supervision: MS, TG; Visualization: RL; Writing – initial draft: RL; Writing – reviewing and editing: RL, AK, MS, SdK, TG.

## Acknowledgements

RL’s work on this project was funded by the Natural Environment Research Council (NERC) EAO Doctoral Training Partnership (grant NE/L002469/1) supported by the Chester Zoo Conservation Scholar and Fellow Scheme. RL also received funding through the JSPS Summer Programme. AK’s work on this project was funded by the Engineering and Physical Sciences Research Council (EPSRC) Doctoral Training Partnership (grant number V520299). This project was also funded by the Daiwa Anglo-Japanese Foundation and JSPS Grants-in-Aid for Young Scientists (grant 23680027 and 16H06177). We thank Nagisa Matsuda, Ken Otsuka, Hiroko Kagawa, and Nao Ota for archival song recordings. We also thank CD Durrant for computational support. Finally, we thank Patrycja Strycharczuk for useful discussions and feedback.

## APPENDIX 1: Results of GLMM examining the effect of song length and note position in songs (with introductory notes removed) on duration. All durations were logged

| **Population level** |  |  |  |
| --- | --- | --- | --- |
| *Question* | *Measurement* | *Song length* | *Position in song (logged)* |
| Are longer songs made up of shorter notes? | Log note duration (log(s)) | -0.003<br>p=0.092 | 0.042<br>p=0.263 |
| Do longer songs contain shorter versions of the same note types? | Deviation from note type mean (log(s)) | 0.00007<br>p=0.850 | 0.0016<br>p=0.949 |
| Are longer songs made up of shorter note types? | Average logged duration of note type (log(s)) | -0.003<br>p=0.003 | 0.038<br>p=0.051 |
| **Individual level** |  |  |  |
| *Question* | *Measurement* | *Mean centred song length* | *Position in song (logged)* |
| Are longer songs made up of shorter notes? | Log note duration (log(s)) | -0.002<br>p=0.084 | 0.037<br>p=0.326 |
| Do longer songs contain shorter versions of the same note types? | Deviation from note type mean (log(s)) | -5.59 x 10<sup>-5</sup><br>p=0.899 | -0.002<br>p=0.943 |
| Are longer songs made up of shorter note types? | Average logged duration of note type (log(s)) | -0.002<br>p=0.013 | 0.033<br>p=0.084 |

